# Stability of dentate gyrus granule cell mossy fiber BDNF protein expression with age and resistance of granule cells to Alzheimer’s disease neuropathology in a mouse model

**DOI:** 10.1101/2023.05.07.539742

**Authors:** Chiara Criscuolo, Elissavet Chartampila, Stephen D. Ginsberg, Helen E. Scharfman

## Abstract

The neurotrophin brain-derived neurotrophic factor (BDNF) is important in development and maintenance of neurons and their plasticity. Hippocampal BDNF has been implicated Alzheimer’s disease (AD) because hippocampal levels in AD patients and AD animal models are consistently downregulated, suggesting that reduced BDNF contributes to AD. However, the location where hippocampal BDNF protein is most highly expressed, the mossy fiber (MF) axons of dentate gyrus (DG) granule cells (GCs), has been understudied, and never in controlled *in vivo* conditions. We examined MF BDNF protein in the Tg2576 mouse model of AD. Tg2576 and wild type (WT) mice of both sexes were examined at 2-3 months of age, when amyloid-β (Aβ) is present in neurons but plaques are absent, and 11-20 months of age, after plaque accumulation. As shown previously, WT mice exhibited high levels of MF BDNF protein. Interestingly, there was no significant decline with age in either genotype or sex. Notably, we found a correlation between MF BDNF protein and GC ΔFosB, a transcription factor that increases after 1-2 weeks of elevated neuronal activity. Remarkably, there was relatively little evidence of Aβ in GCs or the GC layer even at old ages. Results indicate MF BDNF is stable in the Tg2576 mouse, and MF BDNF may remain unchanged due to increased GC neuronal activity, since BDNF expression is well known to be activity-dependent. The resistance of GCs to long-term Aβ accumulation provides an opportunity to understand how to protect other vulnerable neurons from increased Aβ levels and therefore has translational implications.

**SIGNIFICANCE:** Declining hippocampal brain-derived neurotrophic factor (BDNF) has been implicated in the pathogenesis of Alzheimer’s disease (AD). However, few studies have examined where hippocampal BDNF protein has its highest concentration, and plays a critical role in memory, the dentate gyrus granule cell (GC) axons (mossy fibers; MFs). Using a well-established mouse model of cerebral amyloid overexpression, the Tg2576 mouse model of AD, we found that MF BDNF did not decline with age, suggesting a notable exception to the idea that reduced hippocampal BDNF contributes to AD pathobiology. We also identified that Tg2576 GC activity correlates with MF BDNF protein based on GC expression of the transcription factor ΔFosB. These data are consistent with the activity-dependence of MF BDNF. In addition, we found that Tg2576 GCs were relatively resistant to accumulation of amyloid-b, providing insight into AD resilience, which has strong therapeutic implications.

## INTRODUCTION

Brain-derived neurotrophic factor (BDNF) is a member of the neurotrophin family of growth factors (Barde etal 1982, Chao etal 1998, Teng & Hempstead 2004). BDNF is important in neuronal development and is also critical in the adult brain, where it supports neuronal structure and plasticity (Pardon 2010, Park & Poo 2013). One critical brain region where BDNF is high and has been studied extensively is the hippocampus, where it is considered to be instrumental to learning and memory (Yamada & Nabeshima 2004, Minichiello 2009).

AD brain tissues show variable BDNF levels, with some increases and some decreases depending on the brain area and the cell type being investigated. In hippocampus there is not always a significant change in AD (Ferrer etal 1999, Holsinger etal 2000, Garzon etal 2002, Michalski etal 2015), although some studies have reported that BDNF protein and mRNA decline (Phillips etal 1991, Hock etal 2000), and data from CA1 pyramidal neurons show a robust decline across the progression of dementia that correlates with cognitive decline and neuropathology (Ginsberg etal 2006, Mufson etal 2007, Nagahara etal 2009, Ginsberg etal 2010, Ginsberg etal 2019). In animal models of AD, there are both increases and decreases in hippocampal BDNF mRNA and protein levels (Burbach etal 2004, Szapacs etal 2004, Peng etal 2009). Despite the often equivocal findings, many investigators conclude that reduced BDNF occurs in AD, and contributes to it (Peng etal 2009, Xue etal 2022).

There are numerous studies of serum BDNF in AD but serum levels may not be directly related to brain levels. One reason is that BDNF is expressed at high concentrations in platelets. Nevertheless, reduced serum BDNF has been commonly reported in AD (Altar etal 2009, Erickson etal 2010, Allen etal 2011, Thompson Ray etal 2011, Autry & Monteggia 2012, Tanila 2017). Variability may be explained by the stage of AD, because two studies showed elevated serum BDNF early in AD, at the stage of mild cognitive impairment (MCI) (Laske etal 2007, Angelucci etal 2010) followed by a decline (Laske etal 2007). However, others found decreased serum BDNF both in MCI and later (Yu etal 2008, Forlenza etal 2010).

In normal rats and mice, BDNF protein shows abundant expression in the hippocampus in the mossy fiber (MF) axons of the dentate gyrus (DG) granule cells (GCs) (Conner etal 1997, Yan etal 1997, Dieni etal 2012). Despite the high expression in MFs, to our knowledge only one study has examined MF BDNF in AD and that study used patient-derived tissue (Connor etal 1997). The results suggested decreased MF BDNF protein in AD but there was variation in age of the patients, drug history, postmortem delay, and other factors that could affect BDNF expression levels.

In the present study we took advantage of an antibody to BDNF that shows excellent specificity and staining for MF BDNF levels (Kolbeck etal 1999, Dieni etal 2012). We used an established AD mouse model, Tg2576 mice, which is advantageous because there is a slow development of amyloid-β (Aβ) plaques, occurring after 6 months of age (Citron etal 1992, Hsiao etal 1996, Kawarabayashi etal 2001, Jacobsen etal 2006).Therefore we could sample early (pre-plaque, 2-3 months-old) or late (post-plaque, >11 months old) stages.

The results demonstrated that BDNF protein expression was strong in the MFs in Tg2576 mice and there was no detectable age-related decline. We then asked if the reason BDNF expression remains strong over the lifespan could be related to GC neuronal activity, because BDNF expression increases with activity (Tongiorgi etal 2000) and many AD patients and mouse models of AD exhibit increased excitability (Palop & Mucke 2010, Chin & Scharfman 2013, Vossel etal 2017). In this regard the Tg2576 mouse was useful because Tg2576 mice exhibit increased excitability *in vivo* (Bezzina etal 2015, Kam etal 2016) and in GCs *in vitro* (Alcantara-Gonzalez etal 2021). We found that there were high levels of the transcription factor ΔFosB within GCs in Tg2576 mice when MF BDNF expression was relatively high, supporting the idea that increased GC activity promotes BDNF activity-dependent expression and could explain MF BDNF stability.

We then asked if stable GC BDNF expression might confer protection of GCs from Aβ deposition. Indeed, GCs showed remarkably little evidence of Aβ accumulation using several Aβ antibodies, even at 20 months of age. However, adjacent hilar neurons exhibited robust Aβ accumulation, as did hippocampal pyramidal cells. In summary, these data show BDNF protein in GC MFs is stable with age in Tg2576 mice, that there is a relationship to neuronal activity, and the relative resistance of GCs to Aβ accumulation.

## MATERIALS AND METHODS

### 1. Experimental design

The study used the Tg2576 AD mouse model, with comparisons to wild type (WT) littermates. Two ages were chosen. The first age was 2-3 months, an age when mice are adult but have no sign of extracellular plaques. In addition, mice were selected from ages 11-20 months, when plaques are robust. At each age, both sexes and genotypes were included. In some experiments, estrous cycle stage in female mice was estimated by sampling vaginal cytology at the time of death.

Before perfusion, mice were acclimated for at least 24 hours to the laboratory where perfusion would occur. After perfusion, the brain was sectioned and processed using immunohistochemistry to evaluate BDNF protein expression, DG neuronal activity and Aβ immunoreactivity, as described below.

### 2. Animal care and use

The experimental procedures were performed according to National Institutes of Health guidelines and approved by the Institutional Animal Care and Use Committee at the Nathan Kline Institute. Mice were housed in standard mouse cages, with a 12 hr light-dark cycle. Mice had food (Rodent diet 5001; LabDiet) and water ad libitum. During gestation and until weaning, mice were fed chow formulated for breeding (Formulab diet 5008; LabDiet). Mice were weaned at 23–25 days of age and then were fed a standard rodent chow after weaning (Rodent diet 5001, LabDiet). After weaning mice were housed with littermates of the same sex (maximum four mice per cage). For the experimental procedures, two ages were selected: 2-3 months (mean 98.5 ± 5.4 days; range 70-101, *n* = 12) and 11-20 months (mean 451.8 ± 23 days; range 337-611, *n* = 20). Ages in the Results that are in months were calculated by dividing the age in days by 30.3 because the average number of days/month is 30.3.

### 3. Breeding and genotyping

Tg2576 mice express human APP695 with the Swedish (Lys670Arg, Met671Leu) mutations driven by the hamster prion protein promoter (Hsiao etal 1996). They were bred in-house from male heterozygous Tg2576 and female non-transgenic mice (C57BL6/SJL F1 hybrid, Stock# 100012, Jackson Labs). Genotype was determined using an in-house protocol for detecting the APP695 gene.

### 4. Vaginal cytology

All mice were euthanized between 10:00 A.M. and 12:00 P.M. Cycle stage was estimated by assessment of vaginal cytology collected at the time of death. Characterization of vaginal cells was based on Scharfman et al. (Scharfman etal 2008). The cell types were defined as follows: leukocytes (round, small cells), epithelial cells (oval, intermediate-size, nucleated cells), and cornified epithelial cells (multipolar, large, nucleated cells; see Figure 4). In young mice there was a pattern of vaginal cytology consistent with a cyclic pattern (D’Amour etal 2015) and in old animals there was a pattern consistent with a cessation of cyclic estrous cycles, s either a predominance of leukocytes or epithelial/cornified epithelial cells, suggesting a state of persistent diestrus or persistent estrus respectively (Scharfman etal 2008, Scharfman etal 2009). In some mice, there were few cells in the vaginal sample each day. These animals were either old, and had entered reproductive senescence, or were young, and their estrous cycles had not become regular yet. It is also possible that mice were not cycling well due to stress or other factors such as not being housed with cages of males nearby (Whitten, 1956; Scharfman et al., 2009; D’Amour et al., 2015). To quantify cell type and number, the number of visible cells in a field of view of 400x400 µm was manually counted using Fiji ImageJ (Schindelin etal 2012).

### 5. Anatomy

#### A. Perfusion-fixation and sectioning

Mice were deeply anesthetized by isoflurane inhalation (NDC#07-893-1389, Patterson Veterinary) followed by urethane [(Cat#U2500, Sigma-Aldrich), 250 mg/kg; stock solution 250 mg/mL in 0.9% sodium chloride (NaCl; Cat#S9888); intraperitoneal (i.p.)]. The abdominal cavity was opened with surgical scissors, followed by the heart cavity. A 26-gauge needle was inserted into the heart, followed by perfusion with 10 mL saline (0.9% NaCl in double distilled H_2_O, ddH_2_O) using a peristaltic pump (Minipuls 1; Gilson) followed by 30 mL of cold (4°C) 4% paraformaldehyde (PFA; Cat#19210, Electron Microscopy Sciences) in 0.1 M phosphate buffer (PB; note all buffers were pH 7.4). The brains were removed immediately and postfixed in 4% PFA at 4°C. Notably, when tissue was postfixed with 4% PFA overnight, BDNF antibodies performed poorly. Therefore, shorter durations of postfixation in 4 % PFA were tested (1, 2 and 3 hours). Three hours of postfixation was chosen because it optimized BDNF staining by reducing PFA exposure, and maintained tissue integrity during processing better than shorter postfixation times. In addition, tissue integrity was improved by transferring sections with pasteur pipettes that were heated to melt the tip into a curved shape, instead of brushes.

#### B. Sectioning

After postfixation, the brains were washed in 0.1 M PB. Before sectioning, the brains were hemisected. One hemisphere was cut in the coronal plane and the other in the horizontal plane (50μm-thick sections) using a vibratome; (Model# VT1000p, Leica). Similar septotemporal levels were selected and processed together. The sections were 300 μm apart. The sections were collected in 0.1 M PB. Sections that were not used immediately were stored at −20°C in 30% sucrose (Cat#S8501, Sigma-Aldrich) and 30% ethylene glycol (Cat#293237, Sigma-Aldrich) diluted in 0.1 M PB. We detected no difference in sections stored in 0.1 M PB and the storage solution.

#### C. Immunohistochemistry

##### 1. BDNF immunostaining: Fluorescence and Brightfield

BDNF protein was detected with a mouse monoclonal anti-BDNF antibody that has been validated [(Mab#9, Developmental Hybridoma Bank] using either a fluorescence or brightfield protocol. This antibody is similar qualitatively to the one that was used originally in normal rodents that demonstrated robust MF staining (Conner et al., 1997b) and was characterized and validated as a mossy fiber marker (Dieni etal 2012).

For immunofluorescence, free-floating sections were first washed in 0.1 M Tris buffered saline (TBS, 3 washes for 5 min each) and then blocked with 3% Mouse-on-Mouse blocking serum (M.O.M; Cat#MKB-2213, Vector Laboratories) in TBS for 1 hour. Primary antibody was diluted in a solution of 3% bovine serum albumin (BSA; Cat#A7906, Sigma-Aldrich), 2% donkey serum (DS; D9663, Sigma-Aldrich), and 0.3% Triton X-100 (Cat#X-100, Sigma-Aldrich) in TBS to yield a final concentration of 10 µg/ml anti-BDNF. Sections were incubated for 2 nights at 4°C on a rotator (The Belly Dancer, Stovall). For detection, donkey anti–mouse IgG-488 Alexa Fluor– conjugated secondary antibody was used (1:500; Cat#A21202, Invitrogen) in a solution of 2% DS and 0.3% Triton X-100 in TBS. Sections were incubated for 1 hour at room temperature (RT), washed in TBS, and then TB. Washes were 3 for 5 min each. Labeled sections were mounted onto glass slides and coverslipped with fluorescent mounting medium (Citofluor AF1; Cat#17970-25, Electron Microscopy Sciences).

For brightfield microscopy, free-floating sections were first washed in TBS (3 washes for 5 min each) and then treated with 0.25% H_2_O_2_ (Cat#216763, Sigma-Aldrich), in TBS for 3 min. After three washes of 5 min each in TBS, sections were incubated in 0.3% Triton X-100 in TBS for 20 min at RT. Next, sections were blocked in 1% BSA, 5% normal horse serum (NHS; Cat#S-2000, Vector Laboratories), and 1.5% M.O.M in TBS for 1 hour at RT. Primary antibody was diluted in a solution of 1% BSA, 5% NHS, and 0.3% Triton X-100 in TBS to yield a final concentration of 10 µg/ml anti-BDNF. Sections were incubated for 2 nights at 4°C on a rotator. Then sections were incubated in biotinylated horse anti-mouse IgG secondary antibody (1:500; Cat#BA-2000, Vector Laboratories) in 1% BSA, and 0.3% Triton X-100 in TBS for 3 hours at RT, followed by Avidin-Biotin-Complex (ABC; ABC Elite kit; 1:1,000; Cat#PK-6100, Vector Laboratories) in 1% BSA in TBS for 1 hour at RT. Sections were rinsed in TBS and then Tris buffer (TB). Washes were 3 for 5 min each. Next, sections were reacted with 3, 3′-diaminobenzidine (DAB; Cat#DS905, Sigma-Aldrich; 50 mg/100 ml in 0.1 M TB) in 40 μg/ml ammonium chloride (NH_4_Cl; Cat#A4514, Sigma-Aldrich), 2 mg/ml D(+)–glucose (Cat#G5767, Sigma-Aldrich), 10 mM nickel chloride (NiCl_2_; Cat#N6136, Sigma-Aldrich), 3 μg/ml glucose oxidase (Cat#G2133-50KU, Sigma-Aldrich), and then washed (3 times for 5 min each) in TB. Labeled sections were mounted on gelatin-coated slides (1% bovine gelatin; Cat#G9391, Sigma-Aldrich) and dried overnight at RT. On the next day, sections were dehydrated with increasing concentrations of ethanol (70%, 2.5 min; 95%, 2.5 min; 100%, 5 min), cleared in Xylene (4 min; Cat#534056, Sigma-Aldrich), and coverslipped with Permount (Cat#17986-01; Electron Microscopy Sciences).

Sections were examined using an upright microscope (BX61, Olympus) equipped with brightfield and fluorescence detection. Sections were photographed using a digital camera (Model RET 2000R-F-CLR-12, Q-Imaging) and acquired using ImagePro Plus, v.7.0 (Media Cybernetics). For both fluorescence and brightfield protocols, sections from WT and Tg2576 were processed and photographed together, using the same microscope and software settings. Figures were composed in Photoshop (v7.0. Adobe).

##### 2. ΔFosB

Sections from WT and Tg2576 were postfixed for 1 hour in 4% PFA. Free-floating sections were first washed in 0.1 M TB (3 washes for 5 min each) and treated with 1% H_2_O_2_ in TB for 3 min. After three washes in TB (5 min each), sections were incubated in 0.25% Triton X-100 in TB (TrisA) and subsequently in 1% BSA and 0.25% Triton X-100 in TB (TrisB), for 10 min each. Then sections were blocked in 10% normal goat serum (NGS; Cat#S-1000, Vector Laboratories), 1% BSA and 0.25% Triton X-100 in TB for 60 min at RT. The primary antibody, a rabbit monoclonal anti-ΔFosB antibody (Cat#D358R, Cell Signaling), was diluted in TrisB (final concentration, 1:1000). Sections were incubated overnight at 4°C on a rotator. On the following day, sections were rinsed in TrisA and subsequently in TrisB for 10 min each, then incubated in biotinylated goat anti-rabbit IgG secondary antibody (1:500; Cat#BA-1000, Vector Laboratories), diluted in TrisB, for 60 min at RT, followed by 2 rinses of 10 min each in TrisA, then TrisB. Next, sections were incubated in ABC (ABC Elite kit; 1:1,000) diluted in TrisB for 2 hours at RT. Sections were rinsed 3 times in TB (5 min each) reacted with DAB (50 mg/100 ml in 0.1 M TB), and then mounted and coverslipped as for BDNF (described above). Sections were examined, photographed and figures were prepared as for BDNF.

##### 3. Aβ immunostaining

Aβ-immunofluorescence (Aβ-IF) was detected primarily with an antibody to human Aβ (McSA1) which was raised against the N-terminal fragment (residues 1–12; Grant etal 2000). This antibody can detect the soluble and insoluble forms of Aβ, with a specificity for the Aβ protein without labeling the amyloid precursor protein (APP, Grant etal 2000, Billings etal 2005). We adapted a protocol from Kobro-Flatmoen et al. (2016) using free-floating sections. At first, sections were postfixed in 4% PFA to improve tissue integrity during the antigen retrieval procedure. After 3 hours of postfixation, sections were washed in 0.1 M PB (3 washes for 5 min each) and then treated for 3 hours in 0.1 M PB (60°C). All the following washes and dilutions were performed in 0.1 M PB. Sections were incubated for 20 min in 0.5% Triton X-100. Sections were then blocked for 2 hours (5% NGS) and incubated overnight on a rotator at 4°C in primary antiserum (1:1000, mouse monoclonal antibody to Aβ; Cat#MM-015-P, MédiMabs), 3% NGS, and 0.5% Triton X-100.

Two validated, conventionally used Aβ antibodies (Li etal 2017, Perez etal 2019) were used to confirm results with McSA1: a mouse monoclonal antibody to Aβ residues 1-16 (1:1000; clone 6E10; Cat#803001, Biolegend) or a mouse monoclonal antibody to Aβ residues 17-24 (1:1000; clone 4G8; Cat#800708, Biolegend). For all Aβ antibodies, incubation with primary antisera was followed by 2 hours of incubation with secondary antibody (1:350, goat anti-mouse IgG Alexa Fluor 488; Cat#A1101, Invitrogen).

Sections were examined using a fluorescence microscope (BX61, Olympus) and photographed using a digital camera (Model# Infinity 3-6UR, Lumenera) and Infinity 3 software (v. 6.5.6, Lumenara). As for other photography described above, sections from WT and Tg2576 were processed and photographed together, using the same microscope and software settings.

#### D. Thioflavin-S staining

Aβ plaques were detected following the protocol of Roberson et al. (2007). Sections from WT and Tg2576 were mounted on 0.1% gelatin-coated slides, incubated in a thioflavin-S solution (1% thioflavin S, Cat#T1892, Sigma-Aldrich, in ddH_2_O) for 10 min at RT, dehydrated in a graded series of ethanol (80%, 4 min; 95%, 4 min; 100%, 4 min), cleared in Xylene (4 min), and coverslipped as for BDNF (described above). Sections were examined and photographed as described above for BDNF.

#### E. Quantification

##### 1. BDNF

Quantification of BDNF-immunoreactivity (BDNF-ir) was done by first defining three regions of interest (ROIs) in stratum lucidum (SL), the location of the BDNF-rich MF projections. These three ROIs were at the end of the MF plexus where BDNF-ir is highest. The background value was measured from an ROI in SR, where BDNF-ir is relatively weak. BDNF-ir was calculated by subtracting the background value for optical density from the average optical density of the three ROIs in SL. Then MF BNDF-ir f was normalized to the background of the same section. Three dorsal sections were used, ranging from the septal pole to the point in the rostral-caudal axis where the hippocampus begins to descend ventrally. Sections were approximately 300 μm apart. They were used to obtain an average for a given mouse.

##### 2. ΔFosB

Quantification of ΔFosB-immunoreactivity (ΔFosB-ir) was done by first defining an ROI that encircled the upper blade of the GC layer (GCL). The GCL was defined as the location of packed GC somata. The borders with the hilus and inner molecular layer were defined as the location where GCs were no longer adjacent to one another, i.e., they were >1 GC cell body width apart (Bermudez-Hernandez etal 2017).

The level of ΔFosB immunoreactivity (ΔFosB-ir) in the GCL was quantified using ImageJ software (NIH) on images at 40x magnification. Images were thresholded to create a binary overlay in which ΔFosB-positive nuclei were above threshold and the background was below threshold similar to other studies of GCs stained with an antibody to Prox1 (Bermudez-Hernandez etal 2017) or other immunocytochemical studies of c-Fos-ir GCs (Duffy etal 2013, Moretto etal 2017). For a GC to be considered positive its ΔFosB-ir had to be >2x the background. Three dorsal sections were used to calculate a mean for a given mouse.

##### 3. McSA1, 6E10 and 4G8

###### 3.1 CA1

Quantification of Aβ-IF was done by first defining three ROIs within the area CA1 cell layer that included the highest levels of fluorescence. The background value was measured by an ROI in SLM where IF was not detected. Aβ-IF was calculated by subtracting the background from the average of the values of intensity of the three ROIs. Aβ-IF for each section was then normalized [(optical density) – (background optical density]/(background optical density). These measurement of MF optical density have also been used previously for other immoncytochemical studies (Lee etal 2012, Skucas etal 2013).

###### GCs

Aβ-IF in GCs was quantified from an ROI encircling the upper blade of the GCL. The background value was measured by an ROI in SR, where the Aβ-IF is weak. Aβ-IF was calculated by subtracting the background from the average intensity of the ROI for the GCL. Aβ-IF for each section was then normalized, as described above.

###### Hilar cells

Aβ-IF in hilar cells was quantified by manually counts of Aβ-IF cell bodies in the hilus of the DG. A positive hilar cell was defined by choosing a threshold value which was based on the value for a bright cell in the Tg2576 mice. The threshold had to be high enough so that WT hilar cells would not reach threshold. This choice of threshold was straightforward because immunofluorescence of Tg2576 hilar cells was so bright and immunofluorescence of WT hilar cells was so hard to detect (see figures). The hilus was defined as zone 4 of Amaral (1978).

To compare the immunofluorescence of CA1 and the hilus, the highest optical density measurement in the CA1 pyramidal cells was compared to the highest value for hilar cells.

#### F. Statistical analysis

All data are presented as the mean ± standard error of the mean (SEM). Statistical significance was set at p < 0.05. Statistical analyses were performed using GraphPad Prism Software (https://www.graphpad.com/scientific-software/prism/, RRID: SCR_002798).

Parametric tests were used when data fit a normal distribution, determined by the D’Agostino and Pearson or Shapiro-Wilk’s normality tests. An unpaired t-test was used for two groups. A two-way ANOVA or three-way ANOVA followed by Tukey’s multiple comparisons test were used when comparing two or three independent variables. Interactions of factors are not reported in the Results unless they were significant. For data that did not follow a normal distribution, non-parametric tests were selected. The Mann-Whitney *U* test was used for two groups. For multiple groups, a Kruskal Wallis test was used. When there was a significant heterogeneity of variance as determined by the Brown-Forsythe test, data were log transformed. If this transformation resolved the heterogeneity of variance, parametric statistics were used on the transformed data. If it did not, non-parametric statistics were used on the raw data.

## RESULTS

### 1. Modified procedures for BDNF immunohistochemistry

We optimized an immunohistochemical protocol that allowed us to evaluate the expression of BDNF protein of WT and Tg2576 mice. This modification included reducing exposure to PFA, and instrumenting procedures to handle the weakly fixed, and therefore fragile, tissue sections. Hippocampal sections were incubated with a highly specific affinity-purified monoclonal antibody to BDNF (see Methods; Fig. 1). Two different protocols were used, one for immunofluorescence microscopy and the other for DAB brightfield illumination (Fig. 1). The results were similar for both protocols. BDNF protein expression was predominantly in the MFs (Fig. 1A-B). A similar pattern of expression was identified both in coronal (Fig. 1A) and horizontal sections (Fig. 1B).

**Fig. 1.**
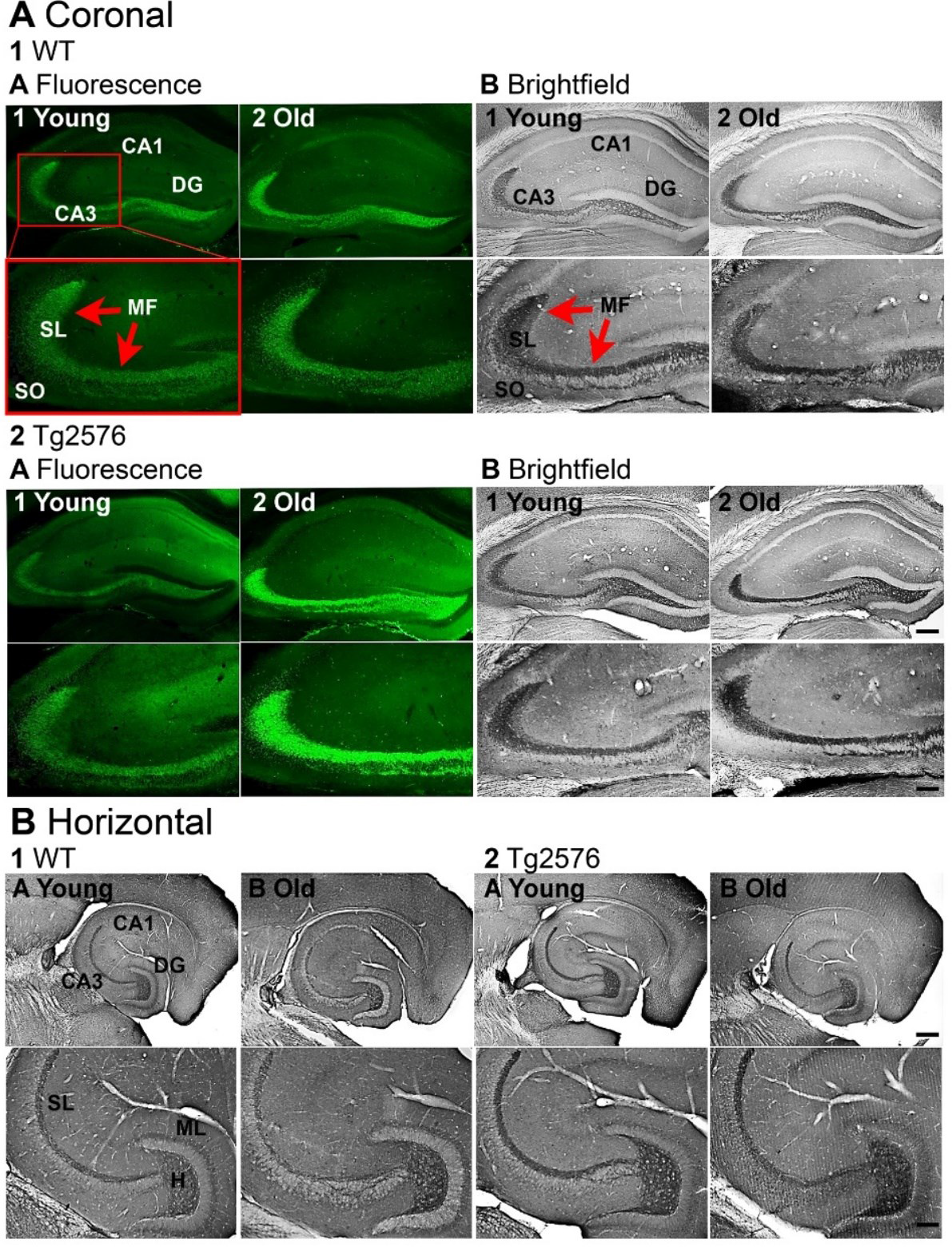
Modified methods allowed visualization of robust BDNF protein expression of in GC mossy fibers (MFs) A. Representative examples of BDNF-immunoreactivity (BDNF-ir) in coronal sections of dorsal hippocampus. Calibration bar is in 2b2 and applies to all images. Top images, 200 µm; Bottom images, 100 µm. SL, stratum lucidum; SO, stratum oriens.

1. WT mice. Intense BDNF-ir was found in MFs (red arrows),

a. Immunofluorescence in a young (1, 3.2 months-old) and old (2, 12.3 months-old) mouse.
b. DAB-ir in a young (1, 3.3 months-old) and old (2, 11.1 months-old) mouse.
2. Tg2576 mice.

a. Immunofluorescence in a young (1, 3.2 months-old) and old (2, 12.2 months-old) mouse.
b. DAB-ir a young (1, 3.3 months-old) and old (2, 10.9 months-old) mouse.
B. Representative examples of BDNF-ir in horizontal sections of ventral hippocampus.

1. WT mice. BDNF-ir was detected in MFs in SL and the hilus (H). Calibration bar is in 2b2. Top, 200 µm; Bottom, 100 µm. ML, molecular layer. a-b. DAB-ir in a young (a, 3.3 months-old) and old (b, 14.2 months-old) mouse.
2. Tg2576 mice. a-b. DAB-ir in a young (a, 3.3 months-old) and old (b, 14.2 months-old) mouse.

### 2. MF BDNF protein is not significantly different in young and old Tg2576 mice

We investigated BDNF-ir in Tg2576 mice and in their WT littermates at two different ages, 2-3 months, and >11 months. Initial studies of mice >11 months was limited to 11-16 months to keep the age range from being too broad. Representative examples of coronal sections from each experimental group are shown in Fig. 2A-B. The groups were young WT, old WT, young Tg2576 and old Tg2576 (n=6/group, 3 males and 3 females).

**Fig. 2.**
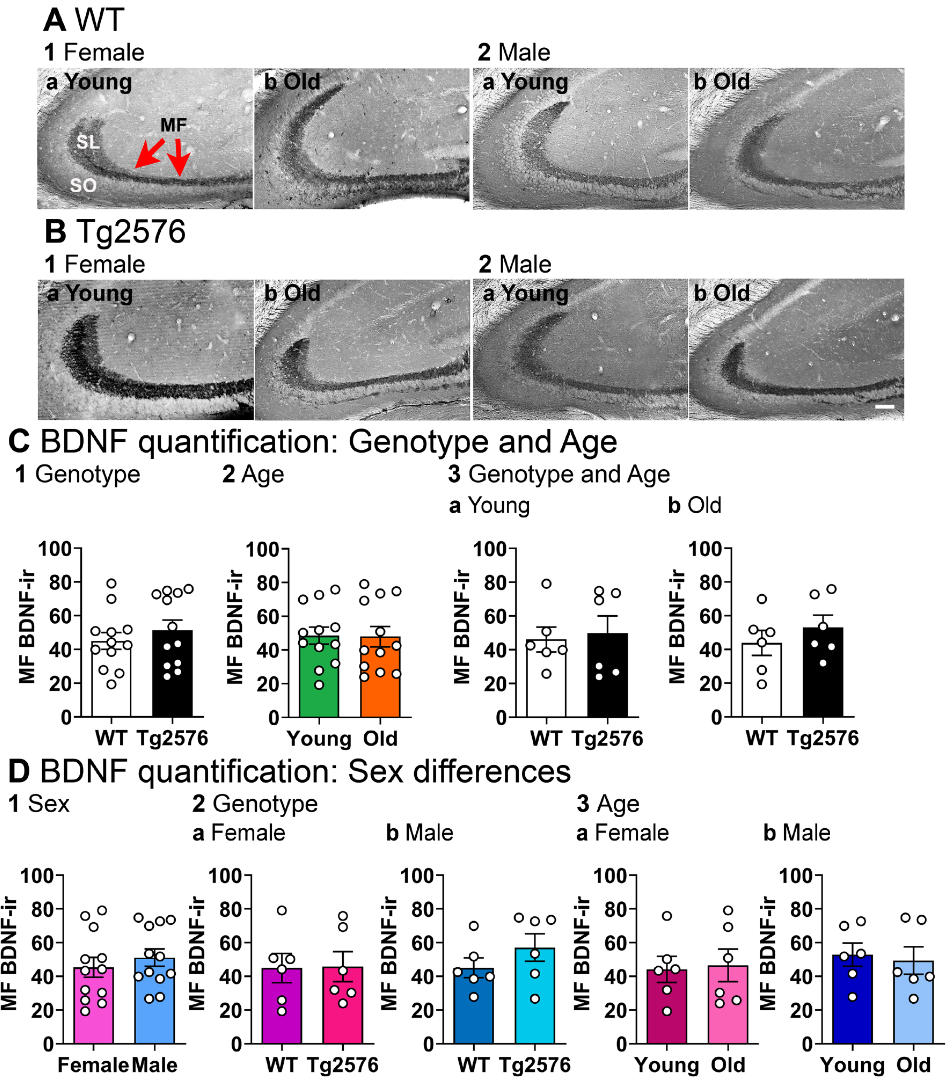
No detectable differences in BDNF protein with age and no sex differences. A. WT mice. Representative examples of MF BDNF-ir (red arrows) in coronal sections of dorsal hippocampus. Calibration bar is in 2b and is 100 µm. SL, stratum lucidum; SO, stratum oriens, MF, mossy fiber.

1. Female mice. A section from a young (a, 2.5 months-old) and old (b, 14.2 months-old) mouse is shown.
2. Male mice. A section from a young (a, 3.2 months-old) and old (b, 12.3 months-old) mouse is shown.
B. Tg2576 mice. Representative examples are shown.

1. Female mice. A section from a young (a, 3.5 months-old) and old (b, 14 months-old) mouse is shown.
2. Male mice. A section from a young (a, 3.3 months-old) and old (b, 12 months-old) mouse is shown.
C. C.

1. A Mann-Whitney *U* test was conducted to compare genotypes with sexes pooled. There was no significant difference in MF BDNF-ir (p = 0.551, *n* = 12/group).
2. when To examine possible differences in ages, genotypes were pooled. There was no significant difference in MF BDNF-ir of young and old mice (Mann-Whitney *U* test, p = 0.712, *n* = 12/group).
3. There was no effect of genotype on MF BDNF-ir at different ages.

a. Young WT and young Tg2576 mice were not different (Mann-Whitney *U* test, p = 0.485, *n* = 6/group).
b. Old WT and old Tg2576 mice were not different (Mann-Whitney *U* test, p = 0.937, old WT vs old Tg2576 *n* = 6/group).
D. D.

1. A Mann-Whitney *U* test was conducted to compare sexes. Genotypes were pooled. There were no significant sex differences (p = 0.514, *n* = 12/group).
2. There was no effect of genotype on MF BDNF-ir in different sexes. Ages were pooled.

a. Female WT and female Tg2576 mice showed no differences(Mann-Whitney *U* test, p = 0.937, *n* = 6/group).
b. Male WT and male Tg2576 mice were not different (Mann-Whitney *U* test, p = 0.240, *n* = 6/group).
3. There was no effect of age on MF BDNF-ir in different sexes.

a. Young female and old female mice were not different (Mann-Whitney *U* test, p = 0.937, *n* = 6/group).
b. Young and old male mice were not different (Mann-Whitney *U* test, p = 0.818, *n* = 6/group).

First we asked if there was a difference in WT and Tg2576 MF BDNF. There were no significant differences in MF BDNF-ir related to genotype when we pooled sexes (Mann-Whitney *U* test, p = 0.551, WT *n* = 12, Tg2576 *n* = 12; Fig. 2C1). There was no significant effect of age when we pooled genotypes (Mann-Whitney *U* test, p = 0.712, young *n* = 12, old *n* = 12; Fig. 2C2). At young ages, WT and Tg2576 MF BDNF-ir were not significantly different (sexes pooled; Mann-Whitney *U* test, p = 0.485, young WT *n* = 6, young Tg2576 *n* = 6; Fig. 2C3a). MF BDNF protein in old Tg2576 mice was not significantly different from old WT mice (Mann-Whitney *U* test, p = 0.937, old WT *n* = 6, old Tg2576 *n* = 6; Fig. 2C3b). These data suggest that MF BDNF-ir did not decline with age.

In Fig. 2D we focused on potential sex differences. There were no significant effects of sex on MF BDNF protein (genotypes pooled, Mann-Whitney *U* test, p = 0.514, female *n* = 12, male *n* = 12; Fig. 2D1). WT females and males were not significantly different (Mann-Whitney *U* test, p = 0.937, n= 6/group), and Tg2576 males and females were not significantly different either (Mann-Whitney *U* test, p = 0.240, *n* = 6/group; Fig. 2D2). There also did not appear to be a decline in MF BDNF-ir with age in females, because there were no significant differences in young and old females (Mann-Whitney *U* test, p = 0.937, young female *n* = 6, old female *n* = 6; Fig. 2D3a) or young and old males (Mann-Whitney *U* test, p = 0.818, young male = 6, old male *n* = 6; Fig. 2D3b).

We then asked if there would be a decline in MF BDNF-ir if older mice were used. Therefore, we used 17-20 months-old mice (WT *n* =4, Tg2576 *n* = 4). We confirmed the results at younger ages, that MF BDNF protein was similar in WT and Tg2576 mice (unpaired t-test, p = 0.457, t = 0.795, df = 6). We also compared the three ages in the two genotypes, pooling sexes. A two-way ANOVA showed there was no effect of age (F(2,26) = 0.00911; p= 0.0991) and a lack of effect of genotype (F(1,26) = 0.820; p = 0.373).

Although these results suggested there were no detectable abnormalities in MF BDNF protein in Tg2576 mice, there were abnormalities in BDNF protein distribution outside the MFs. As shown in Figure 3, we found BDNF-ir in and around extracellular plaques both in hippocampus and cortex (Fig. 3A-B). In these sections, the antibody to BDNF was used and then tissue sections were stained for thioflavin-S to identify plaques (see Methods). The results are consistent with a previous study conducted in the APP23 mouse model of AD, where BDNF mRNA and Congo Red (which can be used as a marker of plaques; (Cohen & Connors 1987)) were double-labeled in cortex (Burbach etal 2004),

**Fig. 3.**
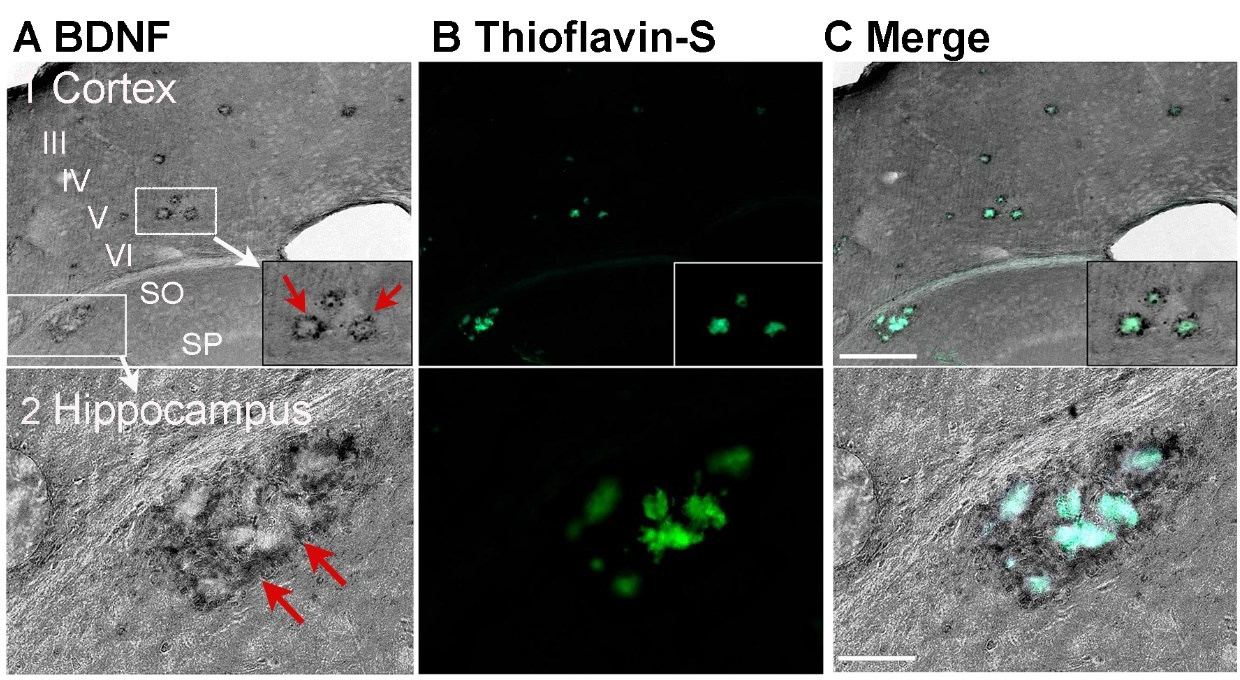
BDNF protein is present around hippocampal and cortical plaques in Tg2576 mice. A. BDNF-ir in a section from a Tg2576 mouse (14.2 months-old) shows BDNF-ir surrounding what appear to be extracellular plaques. Calibration is in C. 1, 200 µm. 2, 100 µm. SO, stratum oriens. SP, stratum pyramidale.

1. Cortex. The cortex above hippocampus is shown. Layers III-VI are marked. There are two boxes that surround extracellular plaques. One box in cortex is expanded at the lower right as marked by the white arrow. It shows BDNF-ir surrounding what appear to be plaques (red arrows).
2. Hippocampus. The second box in A1 is expanded to show the BDNF-ir around plaques in area CA1.
B. The same section was stained for thioflavin-S after immunocytochemistry using the antibody to BDNF. It shows the areas of BDNF-ir are around thioflavin-S staining.
C. A merged image shows the thioflavin-S staining and BDNF-ir overlap. Note that DAB-ir was dark grey and thioflavin-S staining was green, but the merge showed a blue color where there was double-staining.

### 3. MF BDNF expression in female WT and Tg2576 mice is related to estrous cycle phase

The analysis above pooled all female mice, regardless of the stage of the estrous cycle. However, past work has shown that rat MF BDNF rises at as estrogen increases on proestrous morning, remains elevated the next morning (estrous morning) and then returns to baseline during the subsequent days of the estrous cycle, diestrus 1 and diestrus 2 (Becker etal 2005, Scharfman etal 2008, Scharfman etal 2009, D’Amour etal 2015, Li etal 2018).Therefore, we reexamined MF BDNF-ir in a cohort of females where we estimated the cycle stage at the time of perfusion (young *n* = 6, old *n* = 6; WT *n* = 6, Tg2576 *n* = 6). Perfusion was done in the morning (10:00 a.m.-12:00 p.m.). The vaginal sample was taken immediately afterwards. If a vaginal sample had primarily leukocytes, the cycle stage at the time of death was estimated to be diestrous1 or diestrous 2 morning. If epithelial or cornified epithelial cells dominated, the stage was estimated to be proestrous or estrous morning.

The results showed that MF BDNF-ir varied with estimated cycle stage (Fig. 4). Quantification of cells in the vaginal sample showed a significant correlation between the type of cells and the intensity of MF BDNF-ir. When leukocytes were numerous, BDNF-ir was relatively low, and when epithelial or cornified epithelial cells were dominant, BDNF-ir was higher (Pearson’s; R^2^ = 0.579, p = 0.037; Fig. 4B1-2). The results are consistent with an increase in MF BDNF after the proestrous morning surge in estrogen, as shown previously for normal rodents. Interestingly, the correlative data did not show any major differences between genotypes or age (Fig. 4).MF BDNF-ir was weak when cell density was low (Pearson’s; R^2^ = 0.622, p = 0.007; Fig. 4B3). This result could be due to several reasons. First, in 2 months-old females, there may not be a well-established cycle yet. Puberty typically ends at 2 months of age but at the onset of adulthood the estrous cycles may be just starting. Low density in old females may reflect mice entering reproductive senescence. In other mice, there may be low cellularity at a particular cycle stage (Li etal 2018).

**Fig. 4.**
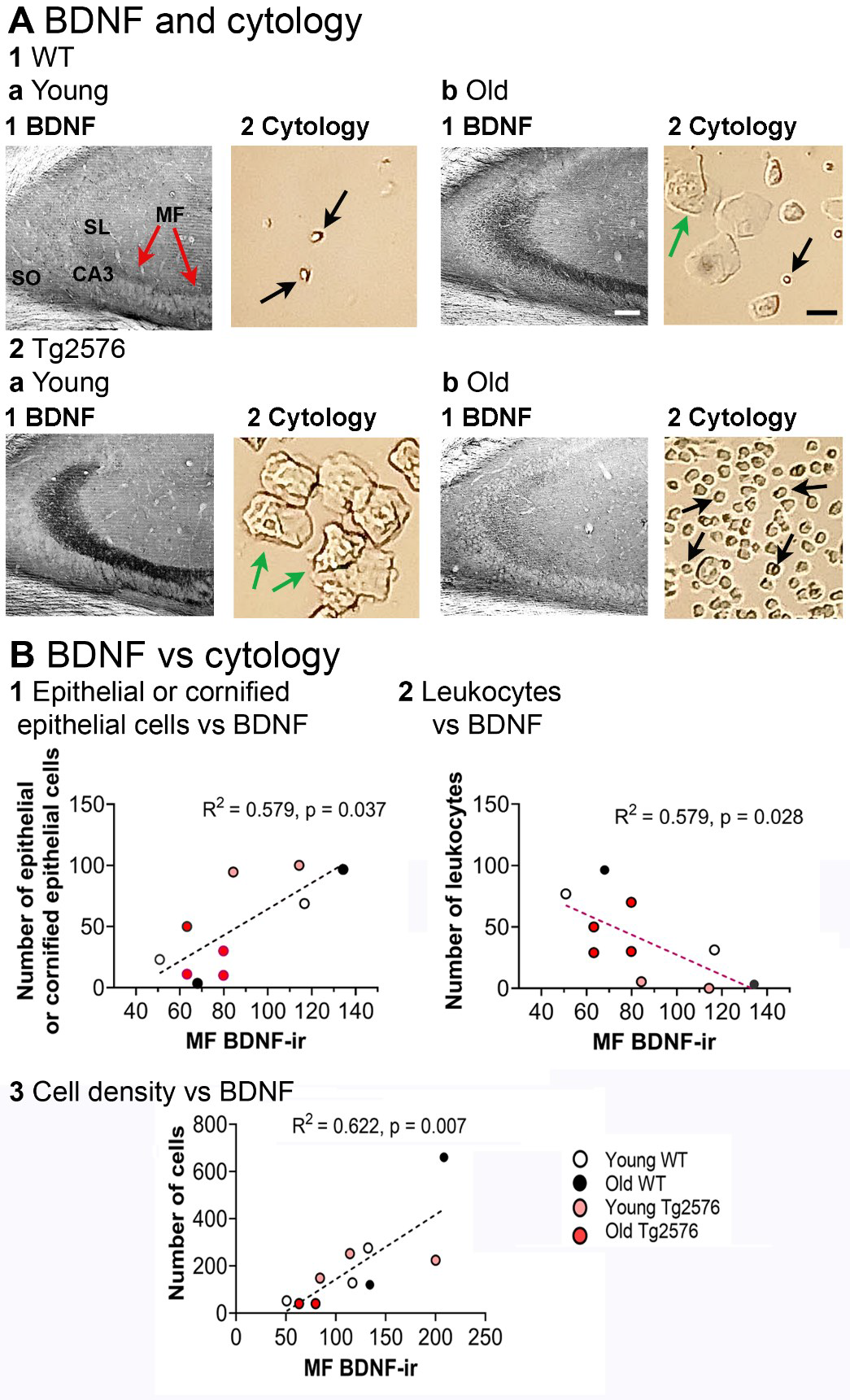
Correlation of MF BDNF protein with estrous cycle phase. A. Examples of BDNF-ir and the vaginal cytology from the same mouse, sampled when the mouse was perfused.

1. WT mice.

a. Young mice. 1-2. BDNF-ir was low when there were predominantly leukocytes in the vaginal sample. An example is from a young WT mouse (2 months-old), but it was found in other experimental groups also. Red arrows point to the MFs. Black arrows point to leukocytes. The calibration bar for all parts of the figure is shown in 1b. For BDNF-ir it corresponds to 100 µm; for Cytology, 25 µm. SL, stratum lucidum; SO, stratum oriens, MF, mossy fiber.
b. Old mice. 1-2. An example of higher BDNF-ir in MFs in a mouse with a vaginal sample that had cornified epithelial cells (green arrow). The example is from an old WT mouse (15 months-old), but it was found in other experimental groups also.
2. Tg2576 mice.

a. Young mice. 1-2. An example of BDNF-ir in a mouse with predominantly cornified epithelial cells. The example is from a young Tg2576 mouse (2 months-old), but it was found in other experimental groups also.
b. Old mice. 1-2. BDNF-ir in a 14.2 months-old mouse with primarily leukocytes in the vaginal sample. This pattern was also found in other experimental groups.
B. BDNF-ir is correlated with cell type and cell density.

1. BDNF-ir was higher when epithelial and cornified epithelial cells were numerous (Pearson’s R^2^ = 0.579, p = 0.037 young, *n* = 6; old, *n* = 6; WT, *n* = 6; Tg2576, *n* = 6).
2. BDNF-ir was lower when leukocytes were numerous (Pearson’s R^2^ = 0.579, p = 0.028 young, *n* = 6; old, *n* = 6; WT, *n* = 6; Tg2576, *n* = 6). These data are consistent with previous studies showing that MF BDNF-ir is lower on diestrous 1 morning in the normal female rat (Scharfman et al., 2003).
3. BDNF-ir was low when the total number of cells was low (Pearson’s R2 = 0.622, p = 0.007). There were 12 mice for this analysis and ages were pooled (young, *n* = 6; old, *n* = 6; WT, *n* = 6; Tg2576, *n* = 6). The data are consistent with the idea that there is lower BDNF when estrous cycles are not robust.

### 5. ΔFosB protein expression in GCs correlates with MF BDNF protein expression

ΔFosB is part of the immediate early gene protein family that are induced by neuronal activation (Minatohara etal 2015). Its long half-life allows it to accumulate in active cells for 1-2 weeks (Nestler etal 2001, McClung etal 2004). In AD, ΔFosB has been shown postmortem and in murine models with mutations in APP that simulate familial AD (Chen etal 1997, Chen etal 2000, Morris etal 2000, Biagini etal 2005, Giordano etal 2015, Corbett etal 2017, You etal 2017, You etal 2018). These data are consistent with the evidence that there is hyperexcitability in AD and in the mouse models (Palop & Mucke 2010, Chin & Scharfman 2013, Vossel etal 2017).

Fig. 5A-B shows ΔFosB-ir in GCs. Consistent with the relative quiescence in GCs normally, WT mice showed the least GC ΔFosB-ir (Fig. 5B2). When a two-way ANOVA was conducted with genotype and age as factors, there was no significant effect of genotype (F(1,16) = 4.143, p = 0.0587) or age (F(1,16) = 0.0908, p = 0.767; Fig. 5C). Similarly, no difference was found when genotype and sex were factors (two-way ANOVA, genotype: F(1,16) = 3.572, p = 0.0770; sex: F(1,16) = 1.251, p = 0.280; Fig. 7E).

**Fig. 5.**
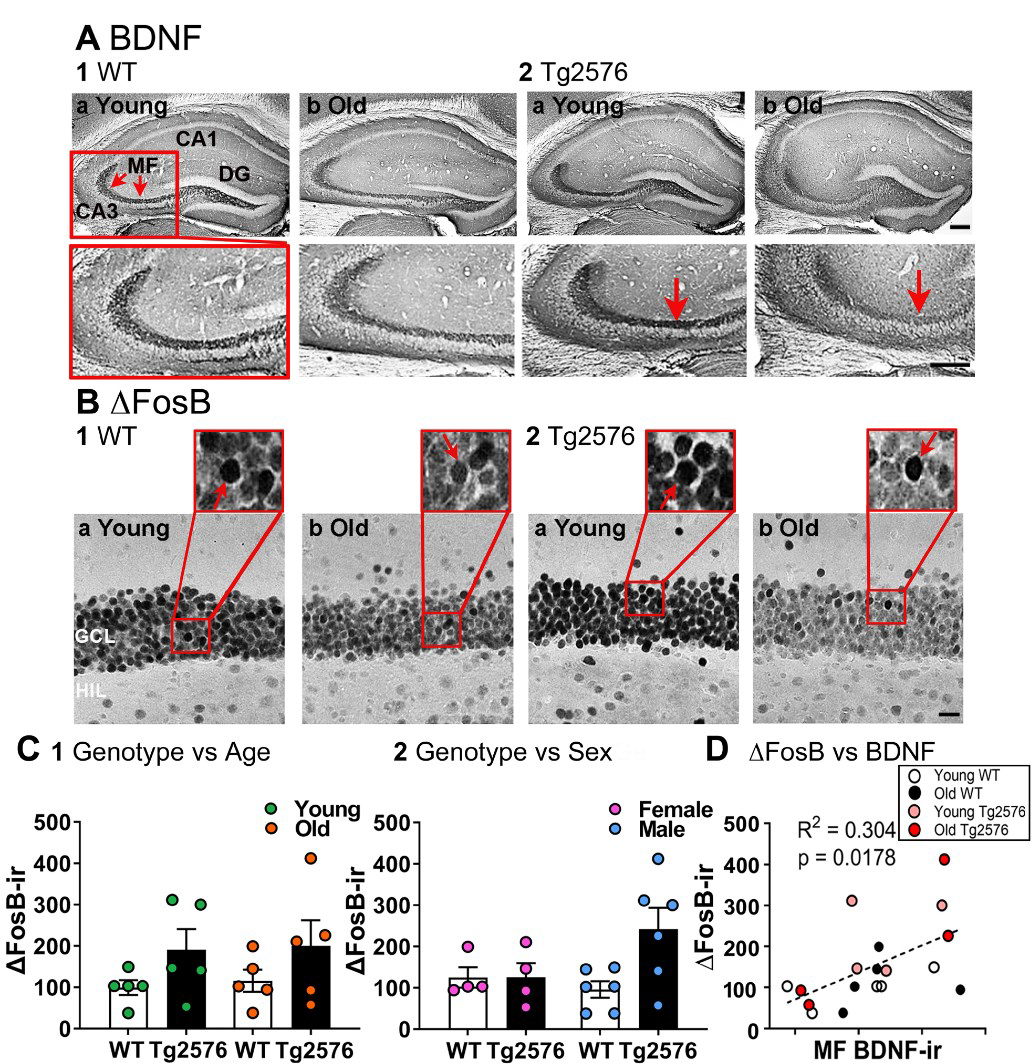
ΔFosB protein is correlated with MF BDNF protein. A. Representative examples of MF BDNF-ir (red arrows) in coronal sections of dorsal hippocampus.

1. WT mice.

a. An example from a young (3.5 months-old) mouse. Red arrows mark BDNF-ir in MFs. The box outlined in red at the top is expanded at the bottom.
b. A section from an old (11.2 months-old) mouse. Note the similarity of BDNF-ir in young and old WT mice.
2. Tg2576mice

a. An example from a young (3.5 months-old) mouse.
b. An example from an old (11.2 months-old) mouse. Note in this example the MFs appear to exhibit less BDNF-ir than the young mouse when one examines area CA3b (red arrow). This example demonstrates that BDNF-ir in Tg2576 mice could appear to decline with age if only CA3b was sampled. This is why MFs were sampled at the end of the terminal plexus near CA2 where BDNF-ir is maximal. Note that the relatively weak staining of CA3b MFs in this Tg2576 mouse was not a consistent finding, another reason it was not the location where BDNF-ir was quantified. Calibration is shown in 2b. Top, 200 µm; Bottom, 100 µm.
B. In the same mice as those shown in A, ΔFosB -ir in the GCL is shown. Insets show ΔFosB-ir is in many GCs (red arrows). 1a-b. Same WT mice as A1a-b. 2a-b. Same Tg2576 mice as A2a-b. Note when BDNF-ir was high in A, ΔFosB -ir was high in B. Conversely, low BDNF-ir occurred when ΔFosB -ir was low. Calibration is in 2b and is20 µm. GCL, granule cell layer; HIL, hilus.
C. Correlation of BDNF-ir and ΔFosB -ir.

1. A two-way ANOVA was conducted with genotype and age as factors. There was no significant effect of age or genotype on ΔFosB-ir (age: F(1,16) = 0.0908, p = 0.767; genotype: F(1,16) = 4.143, p = 0.0587).
2. To analyze sex, a two-way ANOVA was conducted with sex as a factor. There was no effect of sex on ΔFosB-ir (two-way ANOVA, sex: F(1,16) = 1.251, p = 0.280). Genotype was the second main factor, and there was no effect of genotype (F(1,16) = 3.572, p = 0.0770).
D. Higher levels of MF BDNF-ir were correlated with increased ΔFosB-ir in the GCL (Pearson’s R^2^ = 0.304, p = 0.0178, *n* = 20), indicating increased neuronal activity in GCs is related to BDNF protein expression in the GC axons.

Next, we analyzed the relationship between MF BDNF protein and GC ΔFosB-ir. Higher levels of BDNF protein in the MFs were significantly correlated with high GC ΔFosB-ir (Pearson’s, R^2^ = 0.304, p = 0.0178; Fig. 5C). This correlation appeared to be true for both WT and Tg2576 mice (Fig. 5C). Thus, WT BDNF-ir was positively correlated with GC ΔFosB-ir (Pearson’s, R^2^ = 0.567, p = 0.0192; all ages and both sexes pooled). Tg2576 BDNF-ir also correlated with GC ΔFosB-ir (Pearson’s, R^2^ = 0.519, p = 0.0438; all ages and both sexes pooled). The results suggest that when GC activity was high, MF BDNF protein increased independent of genotype and sex.

### 6. Aβ expression in different hippocampal subfields

#### 6.1 Aβ is greater in area CA1 pyramidal cells of Tg2576 mice compared to WT

One would expect greater Aβ in area CA1 pyramidal cells of Tg2576 mice relative to WT and this is what we found (Fig. 6A-B). McSA1 antibody was used to detect Aβ because it detects intracellular (oligomeric) Aβ. The difference between Tg2576 and WT mice was present at both ages 2-3 and 11-17 months (two-way ANOVA, genotype: F(1,20) = 53.170, p < 0.0001; age: F(1,20) = 1.316, p = 0.265; Fig. 6C). Both female and male Tg2576 mice showed significantly more Aβ compared to WT (two-way ANOVA, genotype: F(1,20) = 51.03, p < 0.0001; sex: F(1,20) = 0.509, p = 0.484; Fig. 6D). To determine if sex differences would be significant if analyzed according to age, genotypes were pooled and a two-way ANOVA was performed with age and sex as factors. There were no significant effects (sex: F(1,20) = 0.146, p = 0.706; age: F(1,20) = 0.363, p = 0.553; Fig. 6E). A three way-ANOVA confirmed the effects (genotype: F(1,16)= 47.13, p < 0.0001) but not sex (F(1,16) = 0.470, p = 0.503) or age (F(1,16) = 1.166, p = 0.296).

**Fig. 6.**
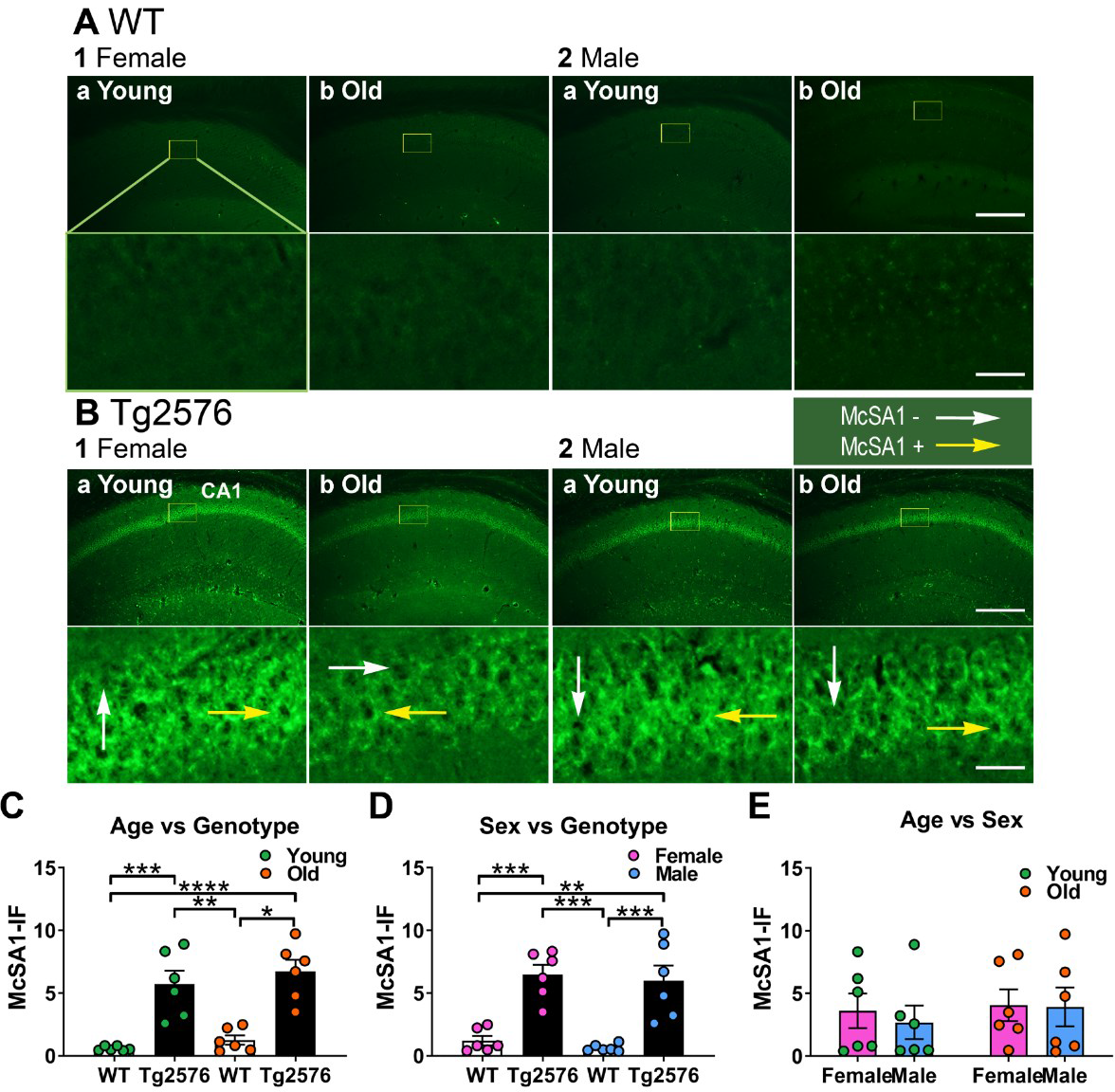
Oligomeric amyloid-β (Aβ) is greater in Tg2576 CA1 than WT in both sexes. A. WT mice. Representative examples of oligomeric Aβ expression in CA1 in coronal sections of dorsal hippocampus revealed by McSA1 staining. As would be predicted in WT mice, low oligomeric Aβ expression was found in all mice.

1. Female mice.

a. Young mice (2.2 months-old). Under the image is an expansion of the area surrounded by a yellow box. Calibration is shown in 2b. Top, 250 µm, Bottom, 25 µm.
b. Old mice (11 months-old).
2. Male mice

a. Young mice (3.4 months-old).
b. Old mice (11 months-old).
B. Tg2576 mice. High levels of oligomeric Aβ expression were found

1. Female mice.

a. A section from a young (3.3 months-old) mouse. Under the image is an expansion of the area surrounded by a yellow box. White arrows indicating McSA1-ir negative cells, yellow arrows indicating McSA1-ir positive cells.
b. A section from an old (15.1 months-old) mouse.
2. Male mice

a. A section from a young (3.2 months-old) mouse.
b. A section from an old (12 months-old) mouse.
C. There was significantly more oligomeric Aβ expression in CA1 of Tg2576 mice compared with WT. This difference was present at young and old ages (two-way ANOVA, genotype: F(1,20) = 53.17, p < 0.0001; age: F(1,20) = 1.316, p = 0.265) followed by Tukey’s multiple comparisons tests (all p < 0.05)
D. In both female and male Tg2576 mice there was a significant difference in oligomeric Aβ compared to WT (two-way ANOVA, genotype: F(1,20) = 51.03, p < 0.0001; sex: F(1,20) = 0.509, p = 0.484) followed by Tukey’s multiple comparisons tests (all p < 0.05).
E. Oligomeric Aβ levels of Tg2576 mice were not dependent on age or sex (genotypes pooled, two-way ANOVA, sex: F(1,20) = 0.146, p = 0.706; age: F(1,20) = 0.363, p = 0.553).

#### 6.2 Surprisingly, GC Aβ expression is similar in Tg2576 and WT mice

We next turned to the DG. Surprisingly, McSA1-IF was not elevated in Tg2576 GCs relative to WT (Fig. 7). When a two-way ANOVA was conducted with genotype and age as factors, there was no significant effect of genotype (F(1,20) = 2.349, p = 0.141) or age (F(1,20) = 0.0186, p = 0.893; Fig. 7D). Similarly, there were no significant effects of genotype or sex (two-way ANOVA, genotype: F(1,20) = 2.541, p=0.127; sex: F(1,20) = 0.111, p = 0.743; Fig. 7E). To investigate age and sex differences further we pooled genotypes and found no effects (two-way ANOVA, age: F(1,20) = 0.0188, p = 0.892; sex: F(1,20) = 0.104, p = 0.751; Fig. 7F). The results were the same by a three-way ANOVA with genotype, age and sex as factors (genotype: F(1,16) = 2.460, p = 0.136; age: F(1,16) = 0.0195, p = 0.891; sex: F(1,16) = 0.0195, p = 0.747; Fig. 7C).

**Fig. 7.**
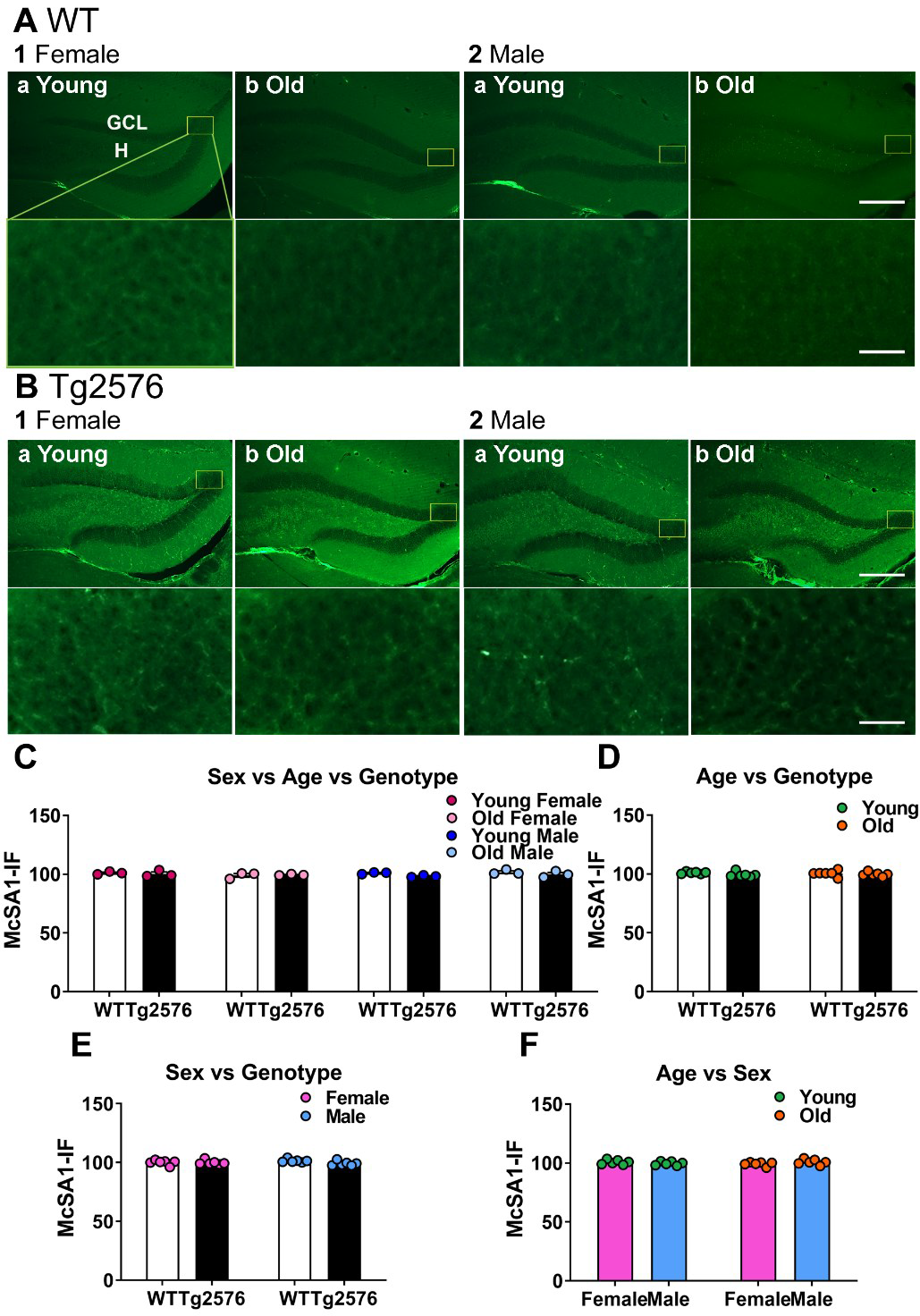
Oligomeric Aβ is low in Tg2576 GCs even at older ages. A. WT mice. Representative examples of McSA1 staining in the GCL of coronal sections from dorsal hippocampus. As one would anticipate from WT mice, expression was hard to detect.

1. Female mice.

a. A section from a young (2.2 months-old) mouse. Under the image is an expansion of the area surrounded by a yellow box. Calibration is shown in 2b. Top, 250 µm, Bottom, 25 µm. GCL, granule cell layer; H, hilus.
b. A section from an old (16 months-old) mouse.
2. Male mice.

a. A section from a young (3.4 months-old) mouse.
b. A section from an old (12.3 months-old) mouse.
B. Tg2576 mice. McSA1 staining was hard to detect in the GCL of all mice.

1. Female mice.

a. A section from a young (2.2 months-old) mouse.
b. A section from an old (14.6 months-old) mouse.
2. Male mice

a. A section from a young (2.3 months-old) mouse.
b. A section from an old (15.1 months-old) mouse.
C. There was no significant effect of genotype, age or sex on McSA1 expression (three-way ANOVA, genotype F(1,16) = 2.465, p = 0.136; age: F(1,16) = 0.0195, p = 0.891; sex: F(1,16) = 0.0195, p = 0.747).
D. There also was no significant effect of genotype or age when sexes were pooled (two-way ANOVA, genotype: F(1,20) = 2.349, p=0.1410; age: F(1,20) = 0.0186, p = 0.892).
E. There was no effect of genotype or sex when ages were pooled (two-way ANOVA, genotype: F(1,20) = 2.541, p=0.127; sex: F(1,20) = 0.111, p = 0.741).
F. There was no effect of age or sex when genotypes were pooled (two-way ANOVA, age: F(1,20) = 0.0188, p = 0.892; sex: F(1,20) = 0.104, p = 0.751).

To validate the use of the McSA1 antibody did not influence the results, we used two additional antibodies to the N-terminal epitope of Aβ (6E10 and 8G10; Fig. 8A-B). The results confirmed that Tg2576 and WT mice were not significantly different (unpaired t-test; 6E10, p = 0.878, t = 0.158, df = 9; WT *n* = 5, Tg2576 *n* = 6; 4G8, p = 0.589, t = 0.563, df = 8; WT *n* = 4, Tg2576 *n* = 6; Fig. 8C-D).

**Fig. 8.**
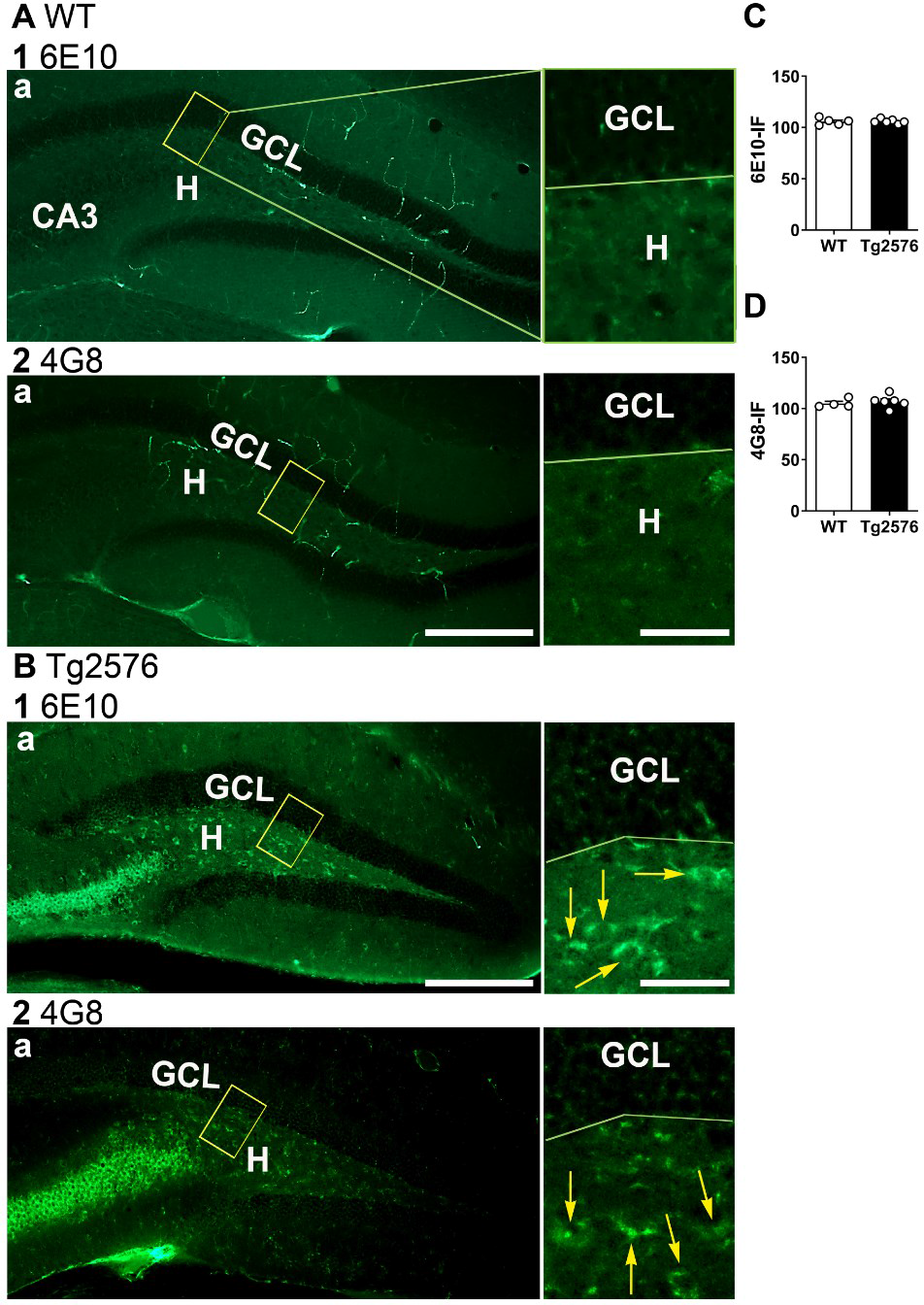
Confirmation of low GC Aβ expression using additional Aβ antibodies. A. WT mice (11 months-old). Representative examples of Aβ-IF in the GCL in coronal sections of dorsal hippocampus.

1. 6E10 antibody.

a-b. Weak staining is shown at low power in a. The yellow box is expanded in b.
2. 4G8 antibody.

a-b. Weak staining is shown at low power in a. The yellow box is expanded in b. Calibration is shown in 2a. Top, 250 µm, Bottom, 25 µm. GCL, granule cell layer; H, hilus.
B. Tg2576 mice (12 months-old). Representative examples of coronal sections of dorsal hippocampus.

1. 6E10 antibody. a-b. Robust staining is shown in CA3 and the hilus but not the GCL.
2. 4G8 antibody. a-b. Strong staining is shown in CA3 and the hilus but not the GCL. Yellow arrows indicate 6E10-ir and 4G8-ir. Calibration is shown in 1a. Top, 250 µm, Bottom, 25 µm.
C. 6E10. There were no significant different in Aβ-IF between WT and Tg2576 mice (sexes pooled; unpaired t-test, p = 0.878, t = 0.158, df = 9; WT *n* = 5, Tg2576 *n* = 6).
D. 4G8. There were no significant different in the Aβ-IF in WT and Tg2576 mice (sexes pooled, unpaired t-test, p = 0.559, t = 0.563, df = 8; WT *n* = 4, Tg2576 *n* = 6).

#### 6.3 Tg2576 hilar cells exhibit robust Aβ although GCs do not

Interestingly, hilar cells showed robust McSA1-IF in Tg2576 mice even though adjacent GCs did not. WT mice did not show any Aβ expression in the hilus (Fig. 9A-B). In Tg2576 mice, there was no significant effect of age on the numbers of McSA1-positive cells (unpaired t-test, p = 0.0545, t = 2.177, df = 10; Fig. 9C) or sex (unpaired t-test, p = 0.235, t = 1.265, df = 10; Fig. 9D).

**Fig. 9.**
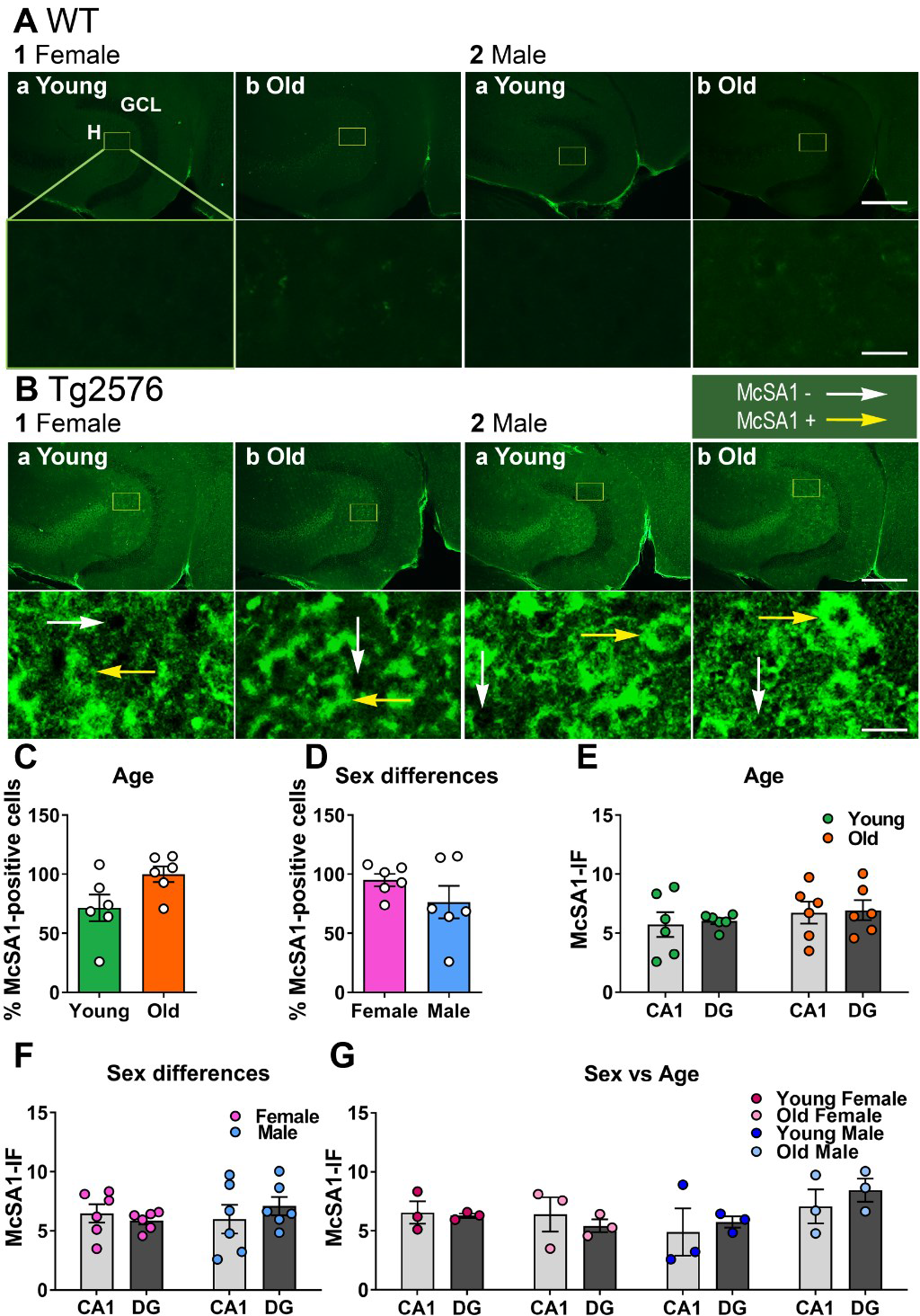
Aβ is elevated in Tg2576 hilar cells. A. WT mice. Representative examples of McSA1 expression in hilar cells in horizontal sections of hippocampus.

1. Female mice. Young (a, 2.2 months-old) and old (b, 16 months-old) examples.
2. Male mice. Young (a, 3.4 months-old) and old (b, 12.3 months-old) examples. Calibration is shown in 1b. Top, 250 µm, Bottom, 25 µm. GCL, granule cell layer; H, hilus.
B. Tg2576 mice. Representative examples of McSA1 expression in hilar cells of horizontal sections of ventral hippocampus. Intense Aβ-IF were found in all mice. White arrows indicate McSA1-negative cells, yellow arrows indicate McSA1-positive cells.

1. Female mice.

a. Young mice (2.2 months-old). The yellow box in the top image is expanded in the lower image. Calibration is shown in 1b. Top, 250 µm, Bottom, 25 µm.
b. Old mice (14.6 months-old).
2. Male mice

a. Young mice (2.3 months-old).
b. Old mice (15.1 months-old).
C. There were no differences in Aβ−IF when young and old Tg2576 mice were compared (unpaired t-test, p = 0.0545, t = 2.177, df = 10). For this comparison sexes were pooled.
D. To examine a sex difference, ages were pooled. There was no sex difference in Aβ−IF of Tg2576 mice (unpaired t-test, p = 0.235, t = 1.265, df = 10).
E. CA1 vs. hilus. A two-way ANOVA with location and age as factors showed no effect of location (F(1,20) = 0.0870, p = 0.771) and no effect of age (F(1,20) = 1.344, p = 0.260).
F. To ask whether one sex might show an effect of location, a two-way ANOVA with location and sex as factors was conducted, and there was no effect of sex (F(1,20) = 0.211, p = 0.651). Similar to E, there was no effect of location (F(1,20) = 0.0868, p = 0.771).
G. To take all factors into account, a three-way ANOVA with location, age and sex as factors was conducted. There were no effects (sex: F(1,16) = 0.213, p = 0.646; age: F(1,16) = 1.394, p = 0.255; location: F(1,16) = 0.0902, p = 0.768).

Because hilar Aβ−IF was so strong (Fig. 9A-B), we compared CA1 and the hilus to determine if hilar cells had the highest Aβ-IF. To this end, the most intense IF in the CA1 cells was compared to the highest intensity for hilar cells, normalized to background in each case. A two-way ANOVA with location (CA1 versus hilus) and age as factors (sexes were pooled) showed no significant differences (location: F(1,20) = 0.0870, p = 0.771; age: F(1,20) = 1.344, p = 0.260; Fig. 9E). When location and sex were main factors (ages were pooled), there were no significant effects either (location: F(1,20) = 0.0868, p = 0.771; sex: F(1,20) = 0.210, p = 0.651; Fig. 9F). There also were no differences by three-way ANOVA, (location: F(1,16) = 0.0902, p = 0.768; age: F(1,16) = 1.394, p = 0.255; sex: F(1,16) = 0.213, p = 0.646; Fig. 9G).

## DISCUSSION

### Summary

This study showed that MF BDNF protein expression levels do not exhibit a significant decline with age in Tg2576 mice. Even when plaques have accumulated, MF BDNF protein remained at a level that was not significantly different from pre-plaque ages. In addition, GCs showed elevated ΔFosB in Tg2576 mice and it was correlated with MF BDNF protein. Potentially for these reasons, GCs showed a resilience to accumulation of intracellular Aβ, despite robust Aβ in adjacent interconnected hippocampal neurons. These findings have profound implication for human AD where vulnerability to amyloid plaques is differentially distributed across the hippocampal formation.

### MF BDNF protein did not decline with age in WT or Tg2576 mice

Our immunohistochemical studies failed to find a significant difference in MF BDNF levels with aging in WT mice or Tg2576 mice. The lack of a decline in BDNF with age in WT mice is consistent with in situ hybridization histochemistry and ELISA in hippocampus of rats where a decline with age was not observed (Croll etal 1998). Other studies of hippocampal BDNF in WT animals come to a similar conclusion (Lapchak etal 1993). However, other studies suggest that BDNF declines with age in humans. The human studies may not be suggesting there is a species difference, however, because the human studies examined circulating levels of BDNF or platelet BDNF concentration, unlike the rodent studies. In humans, BDNF declined with age (Lommatzsch etal 2005, Ziegenhorn etal 2007) and this was associated with hippocampal volume loss (Erickson etal 2010). Therefore, the idea that BDNF declines with age and this predisposes an individual to reduced function is based primarily on serum and platelet-derived BDNF. However, other pools of BDNF, notably those within the DG, may be relatively stable with age.

If BDNF declines in area CA1 or the whole hippocampus in AD, why would that not be the case in the MFs? The resistance of GCs to insult and injury could be a reason (Scharfman 1999). The robust expression of protective BDNF is a possible reason. GC neuropeptide Y (NPY) is also neuroprotective and upregulated by activity, so it also could be involved in GC resilience and/or neuroplasticity (Vezzani etal 1999, Marty 2000). Indeed, NPY is upregulated in GCs in many AD murine models (Palop & Mucke 2010).

Another contributing factor to the stability of MF BDNF could be that GCs are continually being born throughout life and the new neurons would add MF BDNF to the existing MF pathway. Increased neuronal activity increases adult neurogenesis of GCs (Bengzon etal 1997, Parent etal 1997, Parent etal 1998), potentially elevating MF BDNF as new MFs with additional BDNF are added to SL. However, with age the rate of adult neurogenesis declines (Kempermann 2015) and progenitors may become depleted in AD mouse models (Fu etal 2019).

### ΔFosB was elevated in Tg2576 GCs

In AD mouse models there is increased excitability of GCs (Nenov etal 2015, Kim etal 2021, Smith etal 2022) and GCs specifically (Minkeviciene etal 2009, Alcantara-Gonzalez etal 2021). When excitability is sufficiently elevated, seizures occur and the seizures increase GC ΔFosB (You etal 2017, Fu etal 2019, Stephens etal 2020). Consistent with these studies, we found that Tg2576 GCs had elevated ΔFosB. Furthermore, the mice with elevated GC ΔFosB showed higher MF BDNF and the correlation was statistically significant. Although the correlation does not prove that elevated GC activity and ΔFosB led to the increase in MF BDNF, it supports that hypothesis.

### The influence of sex on MF BDNF

It has been shown that as rodents pass through the stages of the estrous cycle, MF BDNF waxes and wanes (Scharfman etal 2003). As females undergo the preovulatory estrogen surge, MF BDNF protein rises, and it remains high until the next morning, estrous morning (Scharfman etal 2003, Scharfman etal 2007, Harte-Hargrove etal 2015). In the present study we found the same appears to be true for both WT and Tg2576 mice.

One of the limitations of our findings is that the stage of the estrous cycle when MF BDNF was examined was only estimated. One vaginal sample was taken at the time of death. One sample is not sufficient to allow one to determine cycle stage definitively because a cyclic pattern can only be ascertained by many consecutive days of assessment. On the other hand, our methods were sufficient to reproduce prior findings that had daily vaginal cytologic examination for several cycles. In other words, mice with higher MF BDNF showed a vaginal cytologic result consistent with proestrous or estrous morning. Therefore, the prediction of estrous cycle stage in the present study appeared to be a good prediction.

### Resistance of GCs to Aβ

A remarkable finding was a relative resistance of GCs to Aβ accumulation. Thus, when sections were processed with an antibody to Aβ that allows one to detect oligomeric (soluble, intracellular) forms of Aβ, as well as amyloid deposited extracellularly (McSA1 or thioflavin-S), staining was negligible in GCs. This result was not because the antibody or thioflavin-S was not able to detect intracellular or extracellular Aβ, because adjacent hilar neurons exhibited robust intracellular Aβ as did CA1. Furthermore, the findings were reproduced with two other antibodies to Aβ.

This result is important because, taken together with the neuroprotective nature of BDNF, GC resistance could be due, at least in part, to their ability to keep producing BDNF. However, the high activity of GCs in Tg2576 mice boosts production of other protective substances such as NPY, as mentioned above. The findings are important because if one can understand the resistance of GCs to Aβ accumulation it might be possible to use that knowledge to delay or prevent Aβ accumulation in vulnerable cells.

### Limitations

One limitation of the study is that the anti-BDNF antibodies could detect the precursor to BDNF, proBDNF, as well as mature BDNF. This is important because the stability of MF BDNF with age could be due to a rise in proBDNF. Another consideration is that we did not study BDNF receptors. Thus, a rise in MF BDNF may not increase BDNF functionally if BDNF receptors, which include full-length TrkB and truncated isoforms of TrkB, decline. There is data in normal rats suggesting that TrkB mRNA declines with age (Croll etal 1998). Other studies suggest that full-length TrkB changes little (Lapchak etal 1993), but truncated isoforms decline (Silhol etal 2005). However, in AD the changes in TrkB may differ from normal aging (Ginsberg et al., 2019a). Indeed, truncated isoforms increase in the 5xAD mouse model rather than declining (Nitzan etal 2022).

## CONCLUSIONS

The results suggest that a decline in BDNF levels does not occur in all of the regions of the hippocampus in the Tg2576 AD mouse model. In particular, the impairments in learning and memory that involve GCs and the MFs are unlikely to be a result of reduced BDNF protein levels. Nevertheless, it is still possible that strategies to increase BDNF, which have been proposed (Nagahara etal 2009, Rahman etal 2023) may be therapeutic, and may be related to activity-dependence, which is still understudied in AD and AD-relevant animal models.

## Acknowledgments

This study was supported by NIH R01 AG908305 and R01 AG077103 and the New York State Office of Mental Health.

